# Taxon sampling unequally affects individual nodes in a phylogenetic tree: consequences for model gene tree construction in SwissTree

**DOI:** 10.1101/181966

**Authors:** Brigitte Boeckmann, David Dylus, Sebastien Moretti, Adrian Altenhoff, Clément-Marie Train, Evgenia Kriventseva, Lydie Bougueleret, Ioannis Xenarios, Eyal Privman, Toni Gabaldon, Christophe Dessimoz

## Abstract

Medium to large phylogenetic gene trees constructed from datasets of different species density and taxonomic range are rarely topologically consistent because of missing phylogenetic signal, non-phylogenetic signal and error. In this study, we first use simulations to show that taxon sampling unequally affects nodes in a gene tree, which likely contributes to controversial conclusions from taxon sampling experiments and contradicting species phylogenies such as for the boreoeutherians. Hence, because it is unlikely that a large gene tree can be reconstructed correctly based on a single optimized dataset, we take a two-step approach for the construction of model gene trees. First, stable and unstable clades are identified by comparing phylogenetic trees inferred from multiple datasets and data types (nucleotide, amino acid, codon) from the same gene family. Subsequently, data subsets are optimized for the analysis of individual uncertain clades. Results are summarized in form of a model tree that illustrates the evolutionary relationship of gene loci. A case study shows how a seemingly complex gene phylogeny becomes increasingly consistent with the reference species tree by attentive taxon sampling and subtree analysis. The procedure is progressively introduced to SwissTree (http://swisstree.vital-it.ch), a resource of high confidence model gene (locus) trees. Finally we demonstrate the usefulness of SwissTree for orthology benchmarking.

## Introduction

Gene tree reconstruction is challenging. Because of the limited amount of information available, gene trees are typically much more difficult to reconstruct than species trees. Gene trees inferred from different datasets of the same family (e.g. nucleotide vs amino-acid sequence, varying taxon sampling) are often topologically discordant [1]. The reasons for analysis artefacts have been studied and discussed extensively, both stochastic (e.g. short sequences, lack of phylogenetic signal) and systematic error (e.g. inappropriate methods or models, insufficient taxon sampling) as well as their combinatorial effects. Methods and models are continuously enhanced, but a sizable fraction of incorrect predictions seems unavoidable. Hence, complementary steps have to be taken to overcome limits. As an example, because evolutionary events other than speciation (e.g. gene duplication, horizontal gene transfer) are rare in most gene families, substantial improvements can be achieved by tree reconstruction methods that take into account a species phylogeny (reviewed in [2]).

SwissTree is a collection of high-confidence model gene trees for the benchmarking of inferred gene relationships. The project was developed within the Quest for Orthologs (QfO) consortium (https://questfororthologs.org), a community effort aiming to improve orthology predictions [3]. Besides SwissTree, two other databases of reference gene trees exist, Orthobench [4] and TreeFam-A [5]. SwissTree is comparatively small and focuses on carefully establishing a reproducible and coherent system for reference gene tree curation. Challenges concern not only tree reconstruction, interpretation, annotation and visualization, but also issues in benchmarking as well as the feasibility of expeditious reactions to requests for taxa that are not part of the current QfO proteome set by the QfO community. Important recent achievements include the construction of a consensus species tree for organisms of the QfO reference datasets (http://swisstree.vital-it.ch/species_tree) [6], which is now used as reference for the interpretation of gene trees in SwissTree. With regard to the QfO benchmarking activities, we studied in detail phylogenomic database concepts to better understand their fundamental different hierarchical levels (e.g. pairwise species comparisons, ortholog groups, hierarchical ortholog groups, reconciled gene trees); such knowledge is important to define suitable benchmarks [7]. Meanwhile, the Orthology Benchmarking Webservice [8] has been developed, which provides - amongst other tests - a comparison of predicted orthologies with those inferred by SwissTree (‘Gold Standard gene tree test’) and the correctness of an orthology-based species tree in comparison to the reference species tree (‘Species tree discordance test’).

With overrepresented clades on the one hand and consecutive long branches on the other hand, the 78-species QfO reference set (1017; http://www.ebi.ac.uk/reference_proteomes; http://swisstree.vital-it.ch/species_tree) is by no means simple. By empirical evidence we assume that any single phylogenetic analysis is unlikely to correctly infer all evolutionary gene relationships of a gene family for all species in the QfO dataset. Thus, we changed strategy and optimized datasets for the prediction of deep divergence patterns in the gene tree as well as for the prediction of individual clades and subclades. In the majority of cases, these phylogenetic gene trees show some contradicting topologies. Tree inconsistency within gene families raises questions of how to distinguish correct from incorrect, how to best summarize and visualize results and how to maintain reference gene trees.

In this study, we first performed a simulation study in which we explored the impact of taxa sampling on individual nodes in a gene tree and compared ways to best summarize results obtained from a set of heterogeneous gene trees. The study leads to an analysis procedure that offers transparency with regard to the final resulting tree, and facilitates the maintenance and extendibility of SwissTree. We apply the approach to a gene family that is difficult to analyze and construct a highly supported model gene tree that is concordant with the reference species tree. Finally, we discuss the perspective of this approach for the construction of sustainable representative gene trees and possibilities for a stepwise automation.

### Conventions. 1

Throughout this article, we use the term ‘clade’ rather than ‘split’ to specify monophyletic groups with at least two members in a gene tree or species tree. This also applies to unrooted gene trees when compared to a rooted - gene or species - tree. 2. When calculating the mean basal aLRT-SH support of a clade in multiple trees, we consider in addition to predictions also missing predictions - which can be deduced from the set of operational taxonomic units (OTUs) - by setting the support value to ‘0’. This measure, which combines quantity and quality of clade predictions, is referred to as ‘mean2’. 3. Gene trees can depict different incidents dependent on the scientific context. Inferred by tree reconstruction methods, gene trees in fact reflect the divergence pattern of individual gene lineages (variants). Often annotated with speciation and duplication events, nodes can have other meanings. For instance in the case of incomplete lineage sorting (ILS), the observed gene tree species tree discordance results from the erroneous interpretation of a node as speciation event rather than the occurrence of a new gene variant. Because SwissTree is generated to benchmark gene relationships (orthology, paralogy, xenology), we design trees to reflect gene locus relations (Fig. 1). The term ‘locus tree’ was introduced by Rasmussen and Kellis for the development of a joint model for phylogenetics and population genetics [9]. Here we use the term ‘gene locus tree’ as opposed to ‘gene lineage trees’; nodes of a gene locus tree represent vertical gene transfer, duplication and lateral gene transfer, nodes of a gene lineage tree depict in addition coalescent effects such as ILS.

**Fig. 1.**
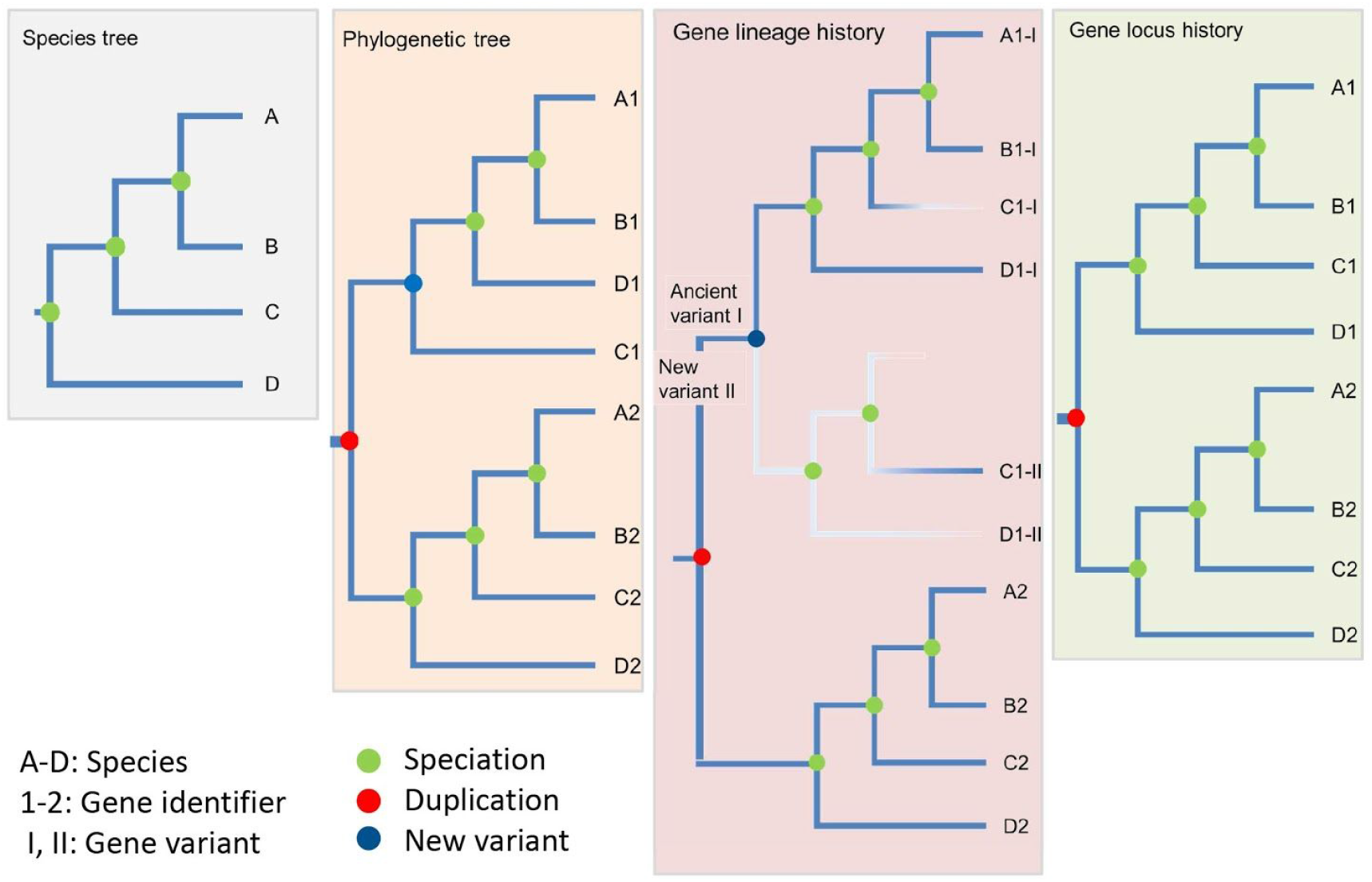
Incomplete lineage sorting: difference between a phylogenetic tree (gene lineage tree) and a gene locus tree. Species tree: evolutionary relationship of species A-D; all nodes in a species tree represent speciation events. Phylogenetic tree: illustration of the evolutionary relationship of gene 1 for species A-D. In the case of incomplete lineage sorting, tree reconstruction methods capture signal originating from retained gene variants that occurred prior to speciation; in this case, the ancestral node represents the occurrence of a new gene variant rather than a speciation event. The difference becomes evident when comparing the gene lineage history and the gene locus history. Gene lineage history: a new gene variant 1-II that appears in the ancestral node is retained in species C and disappears in species D and in the sister clade of C, prior to the speciation of A and B. Simultaneously, the ancient variant is lost in species C, but retained in the other species. The shade of the branch color indicates the variant frequency. Gene locus history: Because gene variants 1-I and 1-II occupy the same gene locus, ILS is not visible in the gene locus tree.

## Results and Discussion

To better understand how different datasets covering the same family contribute to the reconstruction of a gene phylogeny, we first performed a simulation study. While simulation makes many simplifying assumptions, it provides a helpful baseline in which the correct trees are known with certainty. We then show how these results informed the strategy pursued in the construction of reference gene trees in SwissTree. Next, we provide a case study of a family investigated using the SwissTree approach. Based on the same principle, we highlight an efficient way for testing taxa samples for their information content regarding specific clades without tree reconstruction. Finally, we illustrate the usefulness of SwissTree for orthology benchmarking.

### Survey on gene trees inferred from simulated data

To study the problem of gene tree inference from multiple datasets with different taxon sampling, we generate simulation data comprising 100 taxa with 1000 1:1 orthologs that evolve under a codon model with variable rates of sequence evolution across sites and genes. Analyses are performed on the full dataset (“a100”) and six data subsets of 10 and 30 taxa, consisting of two nested subclades (“a10”, “a30”), two balanced sets of taxa (“b10”, “b30”) and two sets of randomly selected taxa (“r10”, “r30”) (for more details, see Material and Methods and Supporting Information S1). Because this study investigates ways to generate representative gene trees from imperfect gene trees, maximum likelihood (ML) tree reconstruction is moderately violated by choosing a fixed model of DNA, codon and amino acid sequence evolution. Possible advantages for codon-based analyses are compensated by a less extensive ML tree search. Gene phylogenies are inferred from the known multiple sequence alignments (MSA), so that erroneous topologies are solely the result of missing phylogenetic signal, non-phylogenetic signal or tree reconstruction artefacts. The final tree set consists of 21,000 trees with 597,000 clades (subtrees of at least 2 OTUs), which we analyze for correctness and support.

The species tree is challenging because it contains long branches and several short internodes (Fig. 2A). First, we explore gene trees that were reconstructed from the known alignment. For the seven datasets (a100, a10-r30) of 1000 gene trees we observe that the fraction of correct tree topologies decreases as the number of genes per tree increases. Remarkably, none of the trees inferred from the full datasets (a100 DNA/codon/aa) is correct (Fig. 2B). By contrast, the fraction of correct clades is primarily linked to the taxon composition with first the balanced subsets, then the complete sets and finally the random subsets. Interestingly, this is less dependent on the size of the dataset. Moreover, most trees possess a small fraction of incorrect clades and only few trees are topologically very distant from the true tree (Fig. 2C). For the 1000 genes from all datasets and data types (21000 trees, 199000 clades), the majority of clades are correctly predicted from all the three data types (DNA, codon, amino acid: 67.64%; 134595 out of 199000 clades); at the other end, 7.23% of the clades (14381) are predicted solely by one of the data types and 13.58% of the clades (27036) are not recovered at all (Fig. 2D). Not surprisingly, the largest fraction of correct clades is predicted from balanced datasets (99.21% for b10, 97.05% for b30), because these have fewer very short (and thus difficult) branches compared with unbalanced trees. The correlation between the level of node support and internode length is shown for dataset a100/aa in Fig. 2E.

**Figure 2.**
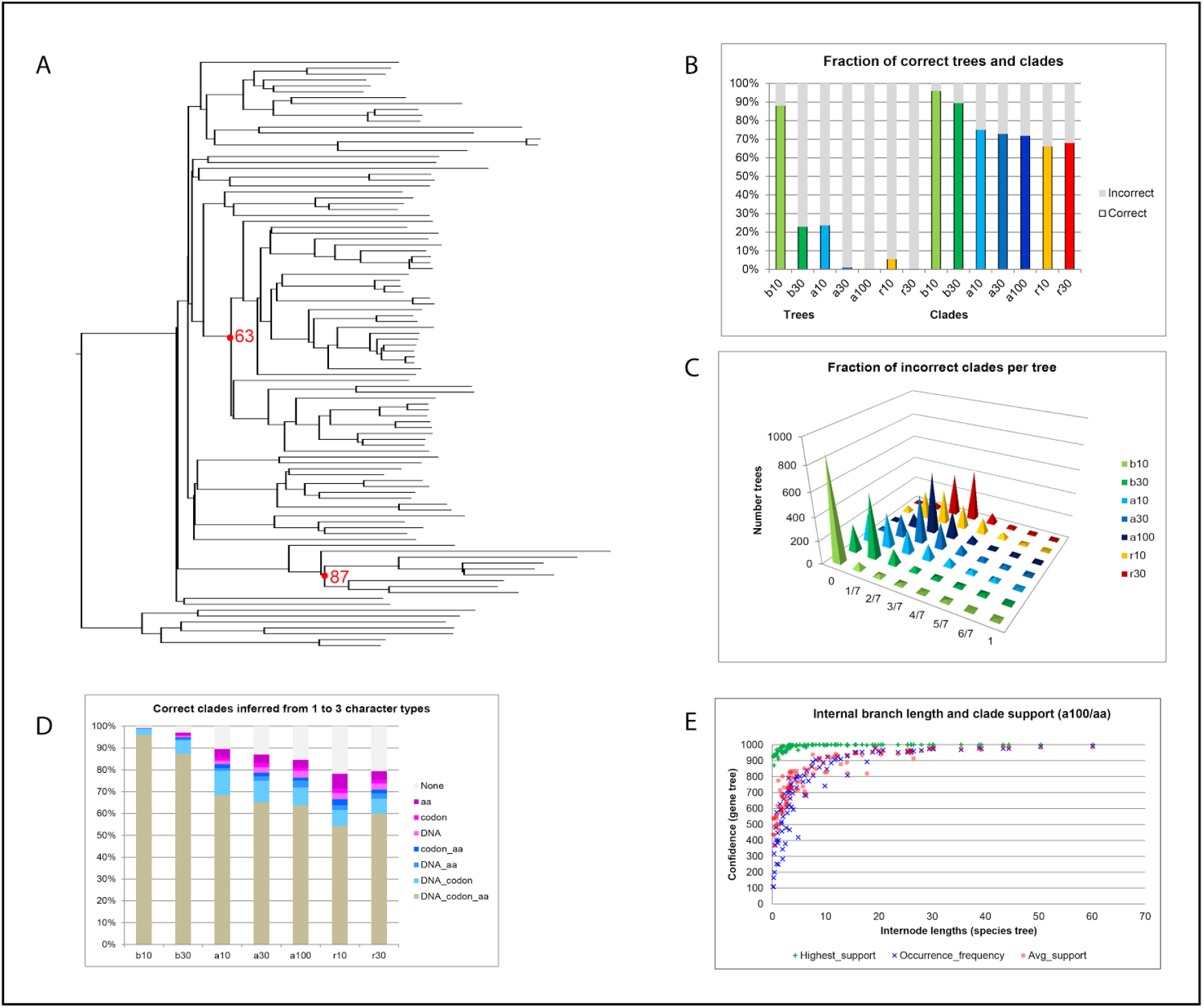
Survey on gene trees inferred from known alignment of simulated data. The simulation data comprises the coding region of 1000 1:1 orthologs from 100 taxa (dataset a100); data subsets consist of two nested subclades of 10 and 30 taxa (a10, a30), two overlapping balanced sets of 10 and 30 taxa (b10, b30) and two sets of 10 and 30 randomly selected taxa (r10, r30). A. Known species tree of the 100 taxa: short internodes and long branches indicate phylogenies that are likely difficult to resolve. Nodes 63 and 78 are annotated. B. Column chart visualizing the correctness of trees inferred from the known MSA of amino acid sequences of the full dataset and the six data subsets. The fraction of correct trees is largest for balanced and small datasets, the fraction of correct clades is approximately consistent within the trait (balanced, all, random) of the taxon composition. Trees inferred from nucleotide or codon sequences show a similar distribution of results with a slightly larger fraction of true trees and true clades. C. Pyramid plot of the average fraction of incorrect clades per tree for the seven protein sequence datasets. D. 100% stacked column chart of correct clades inferred from sequences of all character types (gray; nucleotide, codon, amino acid), by two (blue shade), by one (magenta shade) or not predicted (black). E. Scatter plot showing the branch support of gene trees as a function of the internode length in the species tree (a100/aa). The branch support includes the highest aLRT-SH support for a clade (green plus), arithmetic mean of aLRT-SH support values (blue cross), and clade occurrence frequency (red filled circle).

In summary, the simulated gene phylogenies are difficult to reconstruct because of missing phylogenetic signal and error. None of the largest gene trees is fully correct, but only few trees are topologically very distant from the true tree. The taxon composition of the data subsets has a strong impact on the tree correctness.

#### Clade occurrence frequency and statistical support – which clades are correct?

Since we have established that gene trees are mostly—but not fully—correct, the next question is how can we predict *which* clades are correct. Here we consider three measures: bootstrap branch support [10], aLRT-SH branch support [11], and, since we are particularly interested in situations in which we have multiple datasets covering a single gene family, clade occurrence.

First, we explored the boundary between correct and incorrect clades (“the twilight zone”) in terms of these measures. (Fig. 3). For the 1000 gene trees of each dataset, clade occurrence frequencies are consistently low for incorrect clades and more dispersed for correct clades (Fig. 3A). In contrast, clade occurrence frequencies for incorrect clades show more variation when looking at bootstrap values (20 genes exemplarily selected by increasing alignment length from datasets b30, a30 and r30; see *Materials and Methods*). This is most likely because of biased data and lack of complementary information when compared to results from phylogenomic data (Fig. 3B). As for the aLRT-SH statistics, support is predominantly high for correct clades and dispersed for incorrect clades, though the median support is typically low (Fig. 3C,D). The predictive power of the two measures is thus to some extent complementary. Significant aLRT-SH branch support (>=0.95) is mostly obtained for correct clades; however also a minor fraction of clades (0.000093, 9/97000; a100/aa) show significant support for incorrect predictions in trees that have an average error rate (Supporting Information S2, Fig. S2-1). Thus, a high aLRT-SH is an indication but no guarantee of correctness, nor are strongly supported incorrect clades an indicator of topologically highly erroneous trees.

**Figure 3.**
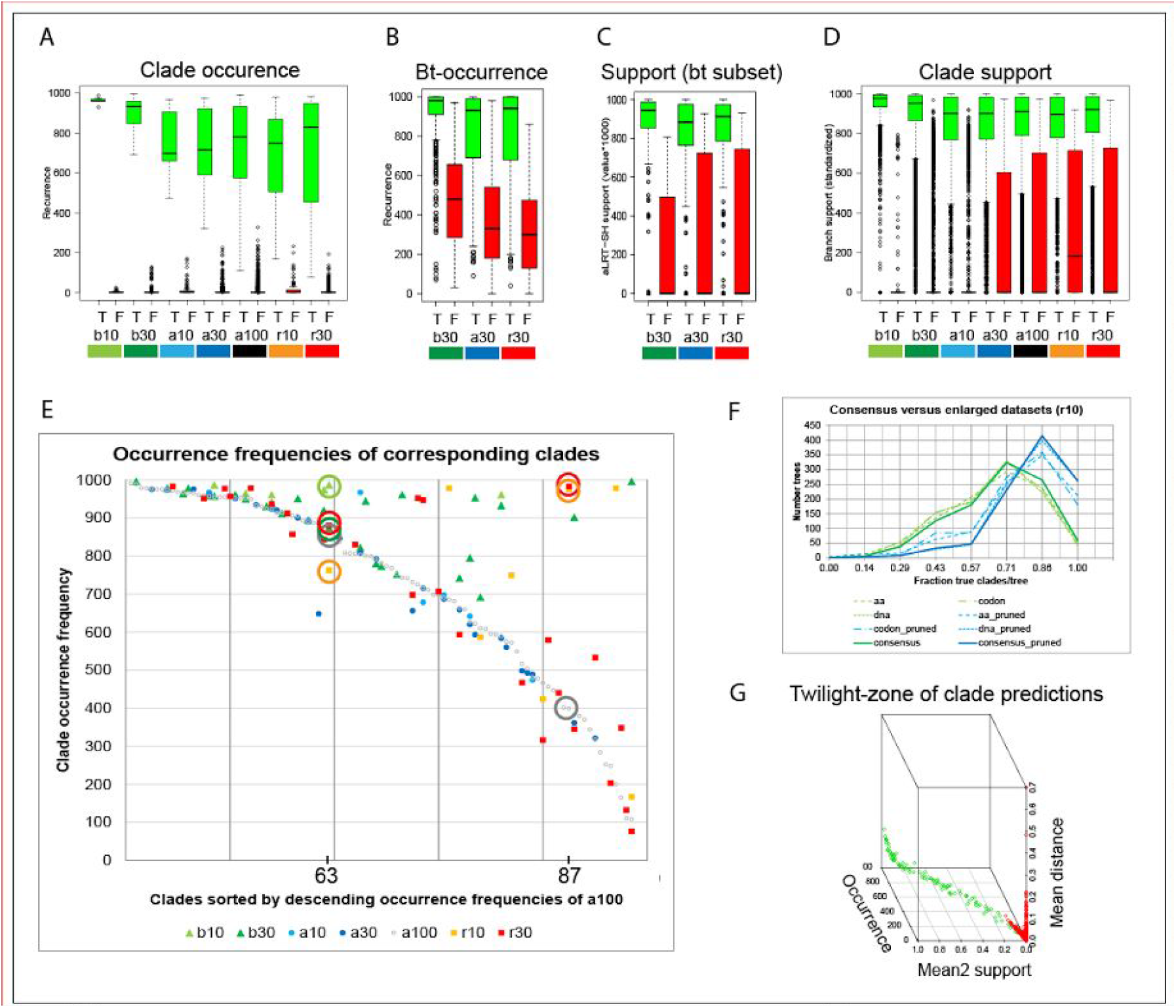
Clade occurrence frequency, branch support and internode length. A-D. Boxplot of clade occurrence frequencies in 1000 genes from dataset a100/aa (A), in trees inferred from 1000 bootstrap replicates for a small data subset of datasets a30/aa, b30/aa and r30/aa. (B), aLRT-SH branch support for the same data as in B (C), aLRT-SH branch support for 1000 genes from dataset a100/aa (D). The box illustrates the Q1-Q3 interquartile range (IQR), the bold line shows the median and whiskers delimit the upper and lower 1.5 IQR; circles depict outliers; support for correct clades is shown in green, support for incorrect clades in red. E. Scatterplot of clade occurrence frequency for corresponding clades of trees from datasets with different taxon composition, sorted by descending support for the dataset a100/aa. Occurrence frequencies of (compatible) clades 63 and 87 from different datasets are marked by circles; colors correspond to the relevant dataset. F. Distribution of the fraction of correct clades per tree for dataset r10 (green dashed lines) and for the corresponding clades from pruned trees of datasets a100 (blue dashed lines); results for the consensus gene trees (DNA/codon/aa) are shown as continuous lines. G. 3D-scatterplot (angle: 300) with clade occurrence frequency (x), mean2 aLRT-SH clade support (y) and the mean basal internode length of each clade (z) (dataset a100/aa).

Clades inferred from datasets with varying taxon composition vary in occurrence frequencies and support (Fig. 3E, Supporting Information S2, Fig. S2-2). For balanced trees (b10, b30), the values for both measures are on average higher than for trees obtained from other datasets, but also trees from random datasets (r10, r30) possess nodes with higher occurrence frequencies and support which can be explained by extended internode lengths due to the lower taxon density. Importantly, this does not mean that smaller datasets generate more accurate phylogenies, which would contradict previous findings ([12], [1]): Fig. 3E also shows that clades with a low occurrence frequency in trees of the full dataset (a100) rarely occur in balanced trees. An easy way to compare tree correctness of large and small datasets is by pruning the larger dataset to the taxa of data subsets. By doing so, the fraction of correct predictions increases for all datasets and data types (dna, codon, aa) with up to 21,3% for trees and up to 15.3% for clades (r10/DNA) (Fig. 3F). Hence, more data generates more accurate tree topologies and multiple datasets a wider spectrum of results. Taken together, concordance between tree topologies from datasets with different taxon sampling constitutes a good indicator for clade correctness.

Further traits of correct topologies are highly supported clades with long basal branch length. Occasionally, an incorrect clade shows similar features (Supporting Information S2, Fig. S2-3), but it is usually not stable and can be easily distinguished from correct clades when summarizing results from multiple analyses (Fig. 3G). Thus, the twilight zone for clade predictions is characterized by rather low - but not lowest - occurrence frequencies and short basal internode distances.

In summary, different taxon sampling have different strengths and weaknesses when it comes to resolve individual nodes. Therefore, it is worthwhile considering multiple datasets of the same gene family when inferring a gene phylogeny.

#### Can we predict correct clades from the sequence alignment?

Phylogenetic trees are typically inferred from MSAs, which means, the alignment contains the phylogenetic information. In cladistics, shared derived characters (synapomorphies, phylogenetic signal) are used to infer species relationships, and in comparative genomics, assumed shared derived (‘synapomorphic’) characters (e.g. nucleotides, codons, amino acids) can be read directly from the MSA for each clade of a tree. However, phylogenetic tree reconstruction methods also have to deal with contradicting (non-phylogenetic) signal, which are signals supporting topologies other than the correct tree. If there exists a correlation between clade occurrence and the fraction of synapomorphic signal, we should be able to efficiently predict suitable taxa samples prior to tree reconstruction.

A simple approach to quantify phylogenetic signal is to determine from the 1000 MSAs the fraction of synapomorphic characters. To further ease the analysis, we focus on individual internodes in the known species tree and determine synapomorphic positions for the known correct tree topology as well as for the two alternative tree topologies around the selected internodes (Fig. 4A). Thereby, we distinguish three modes ‘stringent’, ‘medium’ and ‘relaxed’, which differ in the level of conserved positions within the three subclades and the outgroup (Fig. 4B). Exemplarily, two clades with different occurrence frequencies are chosen from Fig. 3E. The first example, clade 87, is predicted 399 times from dataset a100, but compatible clades are inferred 975 and 985 times from datasets r10 and r30, respectively. The search for synapomorphic sites in the MSAs reveals that there is on average more synapomorphic signal per gene (r10: 11.2, r30: 6.1) and less contradicting signal per gene (r10: 1.7, r30: 0.2) in the random datasets than for the full dataset (a100; synapomorphic: 2.2, contradicting: 1.9) (Fig. 4C). We observe a clear correlation between the fraction of synapomorphic signal (synapomorphic / synapomorphic + contradicting) and the clade occurrence frequency. The second example, clade 63, is inferred more than 750 times from the proteomes of datasets a100 (858), b10 (987), b30 (874), r10 (762), and r30 (880). Nevertheless, the number of synapomorphic signal per gene differs largely for the different datasets and decreases in the order b10 (11.4), r10 (5.1), r30 (1.3), b30 (1.1), a100 (0.4) (Fig. 4D). For dataset a100 the number of signal is below one per gene. When signal is weak, a more fine-grained approach is needed, taking into account amino acid similarity rather than identity.

**Figure 4.**
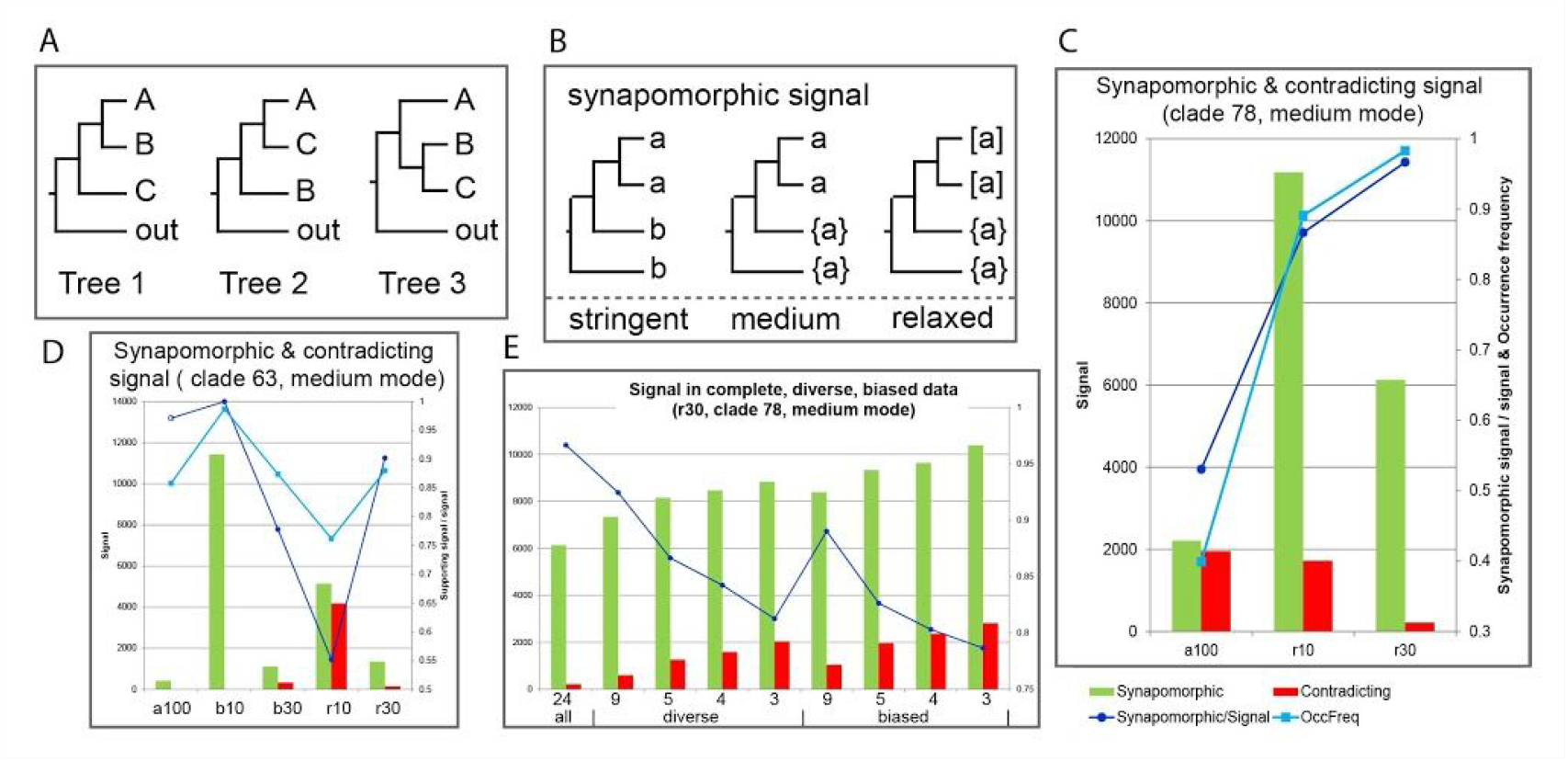
Quantitative assessment of synapomorphic characters in the MSA for tree topologies at two short internodes. A. Correct tree topology (Tree 1) and two alternative tree topologies (Tree 2, Tree 3) around internodes; letters denote clades, ‘out’ stands for outgroup. B. Synapomorphic positions are determined in three modes which are illustrated by trees; stringent: the sister clades and the two outgroups share a common character; medium: the sister clades share a common character that is not present in the outgroups; relaxed: at least one member of each clade of the sister clade shares a common amino acid, that is not present in the outgroups; letters denote amino acids, curly brackets indicate absence of a specific amino acid, square brackets signal presence of a specific amino acid in at least one member of each sister clade; C, D. Column chart of synapomorphic and contradicting MSA positions for clades 87 (C) and 63 (D), and scatter marking in dark blue the fraction of synapomorphic signal (synapomorphic signal / synapomorphic signal + contradicting signal) and in cyan the clade occurrence frequency (OccFreq; clade occurrence în 1000 gene trees). The marker of the fraction for synapomorphic signal is not filled as to indicate a dubious result (less than one signal per gene). E. Column chart of synapomorphic and contradicting MSA positions for clade 87 from dataset r30 with a varying number of members in the largest clade C, which is the outgroup to the true sister clade. See Figure C for the legend; ‘all’ indicates the complete set of clade members (24), ‘diverse’ a taxonomic diverse set of members, ‘biased’ a biased set of more closely related clade members; numbers (x-axis) indicate the number of clade members in a set.

Finally we investigate in a comparison of signal obtained from a complete clade and subclades. We do this for the largest clade in in our example, which is the outgroup of the true sister clade at node 78. Clade subsets with taxonomically diverse clade members provide less signal than a biased subset of clade members (Fig. 4E). In addition we observe that the number of signal increases as the number of clade members decreases. In both cases, the increase of signal is associated with a decrease in the proportion of synapomorphic signal, indicating that biased and small datasets include more noise and less true signal than more balanced and larger datasets.

Three facts are notable in this analysis: 1) the existence of contradicting signal, which has also been found in real data and for which different reasons have been described [13]; 2) the correlation between the fraction of synapomorphic signal and the clade occurrence frequency; 3) the impact of the clade size and taxa diversity on the balance of synapomorphic and contradicting signal. In multiple further examples not described here, we notice that data bias and clade size have a large impact not only on the fraction of synapomorphic and contradicting signal, but likewise on the inferred tree topologies; a large-scale real data example is shown further below. At the comparative genomics level, the investigation of signal in MSAs could therefore constitute a simple and efficient way for compiling genetically diverse, complementing datasets prior to tree reconstruction.

### How to integrate tree inconsistency?

Given a set of almost concordant gene trees, an automated construction of a reference gene tree is feasible. Consensus gene trees, for instance, can be generated from trees of identical taxon composition inferred from the three data types (DNA, codon, aa). By doing so for the simulated data, we obtain only a negligible increase of correct clades, probably due to a lack of complementing signal in the coding sequence (Fig. 3H, Supporting Information S2, Fig. S2-4). When summarizing results from trees for six data subsets including the corresponding three (better) pruned trees of the full dataset, we even observe a decrease of clade correctness over the best gene trees. Another approach could be the construction of supertrees that combine phylogenetic trees of different size and taxon composition and moreover visualizes alternative topologies in form of a network within a tree structure. But in the end, would it really be the gene history that SwissTree strives to reflect? Tree reconstruction methods perceive evolutionary traces of successful gene lineages (variants, alleles). One reason for the often observed gene tree species tree discordance is incomplete lineage sorting (ILS), in which case a gene variant emerges prior to a speciation event, so that the basal node of the relevant clade in fact corresponds to the emergence of a new variant rather than to a speciation event (Fig. 1). In the end, it is a matter of definition on which level gene histories are revealed, interpreted and annotated. Because variants are irrelevant to the attribution of gene relationships, trees in SwissTree present gene histories at the level of gene loci and hence illustrate speciation and changes in gene copy numbers in genomes. Strictly speaking, SwissTree trees are gene locus trees. Notably, this has no impact on the benchmarking of gene relationships with SwissTree, because the tree discordance involves neither gene duplication nor horizontal gene transfer.

Gene locus trees are in principle simpler than gene lineage trees. Nevertheless, the construction of gene locus trees implies distinguishing locus events from incidences of gene lineage events and error. Because species relationships inferred from multiple genes - and a large number of characters - are likely more accurate than when inferred from a single gene, we trust the reference species tree topology more than the topology obtained from a single gene; with other words, we use the species tree model as a null hypothesis that is challenged by discordant findings. As a guideline we presume that tree topologies based on true signal are more robust than analysis artefacts, if the distinct datasets are diverse and compiled for the analysis of a specific node and, moreover, the selected genes evolve at similar rates. In the case of tree inconsistency, if one out of alternative topologies supports the species tree, the tree topology concordant with the species tree gives the more parsimonious explanation unless there is strong evidence for another event but speciation (Fig. 5A). The same applies when datasets lack signal. In the case of consistently discordant topologies or solely discordant topologies, further evidences are looked for, for instance in the MSA, in gene synteny tables and in enlarged datasets (witness of non-orthology, [14]). Thereby, hybridization is difficult to investigate, because it requires a phylogenomic approach and furthermore knowledge on the degree of divergence of the involved species. Nevertheless, it can be considered (and annotated in the reference species tree), if species relationships are affected by hybridization. In the absence of incidences supporting events relevant to the gene locus and the lack of strong support for the dubious topology, we assume incomplete lineage sorting and present the topology that is concordant with the species tree (Fig. 5B). This way, we prevent overprediction of gene locus-relevant events that are due to a lack of signal or erroneous topologies. The benefit of this concept can be tested for the simulation data by mapping identical and compatible subclades from 21 gene trees (3 character types, 7 datasets) to the model of gene locus evolution for the full dataset (a100), which in our case is identical to the species tree. In doing so, the fraction of correctly supported clades increases by 18.37% (13.84% complete clades, 4.53% compatible clades) from 72.04% as achieved from the individual analysis of the a100/aa dataset (a100/aa, 1000 genes, 97000 clades) to 90.41% in the model gene trees - and this without adding additional taxa or clade-specific data optimization (Fig. 5C). We can therefore assume that this approach has the potential for the accurate construction of large gene trees.

**Figure 5.**
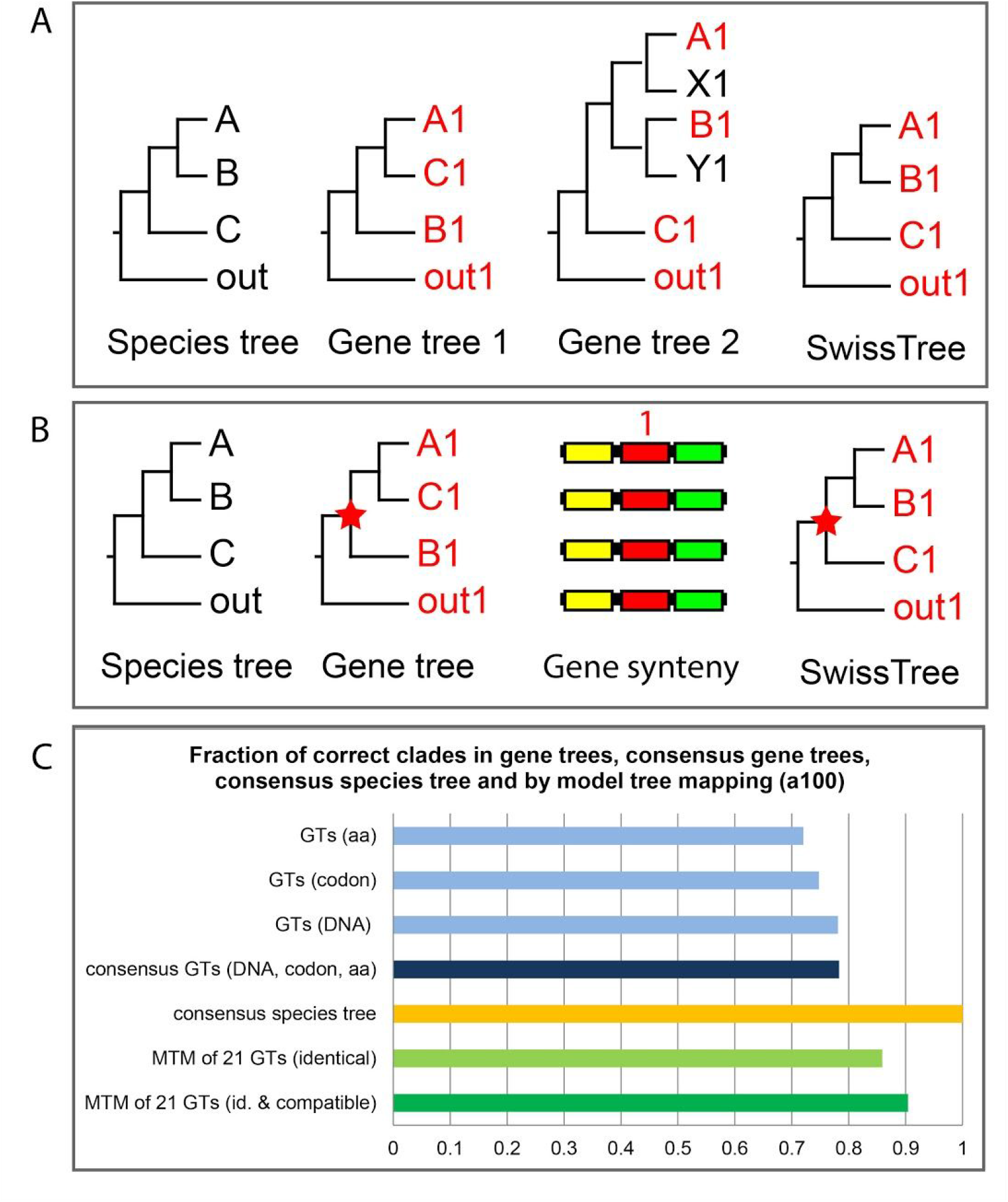
Integration of tree inconsistency. A. Tree inconsistency: if one of the alternative tree topologies supports the species tree, the SwissTree model gene tree corresponds to the concordant tree topology (gene tree 2) for this clade. Alternative topologies are retained in the set of result trees. B. Incomplete lineage sorting (ILS): Trees in SwissTree illustrate gene locus relationships rather than gene genealogies; thus, ILS is not presented in the SwissTree topology. Gene synteny and the lack of gene duplications in related species evidence ILS rather than pseudo-orthology. The red asterisk denotes the polymorphic ancestor. C. Bar chart illustrating clade correctness of gene trees (GTs) for the different data types (amino acid, codon, DNA), of consensus of the gene trees derived from the three different data types (amino acid, codon, DNA), of the consensus species tree from 1000 gene trees, as well as the fraction of correctly reconstructed identical and compatible clades in 21 gene trees identified by model tree mapping (MTM).

### Strategy for the construction and maintenance of representative gene trees

Based on the preceding simulation study we conclude that medium to large phylogenetic gene trees constructed from datasets of different species density and taxonomic range are rarely topologically consistent (Fig. 2B). Nevertheless, we can presume that the majority of clades are predicted correctly (Fig. 2C), because the species diversity - OTU composition and OTU density in the tree space - has a large impact on the level of clade correctness and clade support (Fig. 2B, 3A,D,H) and datasets of complementing taxon composition provide a wide range of results (Fig. 3F,G), topological consistency of compatible clades evidence tree correctness. This finding is supported by MSA analysis for synapomorphic signal: datasets possess different levels of synapomorphic signal for different clades (Fig. 4C,D) and representative datasets of different size possess similar signal for the same clade (Fig. 8; large-scale study below). A second strong indicator of tree correctness is the topological concordance with the species tree and an average high clade support (mean2) (Fig. 3J). As for consensus gene trees, there is a risk of biased tree topologies due to predominantly overlapping signal and a lack of complementing signal in the coding region of the simulated data (Fig. 3B, 5A). As a consequence, trees for SwissTree are generated from multiple datasets - optimized for the analysis of the full dataset, data subsets as well as individual clades and subclades - in a process of repeated analysis, interpretation, and annotation. Results are stored in form of a model gene tree and a tree pool. Figure 6 summarizes the process.

#### Initiation of a tree pool

At first we explore the consistency of tree topologies. A first analysis with (almost) all family members can result in gene trees that are in large part concordant with the species tree, even when inferred from unrevised data. The relevant MSAs typically possess regions of unequal confidence including long indels as well as gaps and missing data that are difficult to interpret by most tree reconstruction algorithms. For the generation of trees for SwissTree, we reassess gene phylogenies using smaller, revised datasets and data subsets for individual clades. Consistent tree topologies inferred from (unbiased) MSAs of different species density and composition evidence phylogenetic signal. In case of alternative topologies, the one concordant with the species tree is the most parsimonious explanation for a gene’s evolution. Alternative topologies that occur with similar frequency indicate a lack of phylogenetic signal. With regard to the statistical branch support, we showed in the simulation study that values can be strongly influenced by the internode length (Fig. 2E). For SwissTree we currently use bootstrap, a rather conservative measure. When trees from extended datasets are pruned, we think it is admissible to report the highest support for a clade of interest; though, we have not yet introduced this in practice. If all phylogenies for a clade are discordant from the species tree, we perform additional studies that focus on individual spots in the tree and strive to explain topologies, taking into account also gene synteny, if species are not too distant from each other. Finally, representative trees - including the ones with contradicting topologies - are gathered in the family ‘tree pool’.

#### Model gene trees and confidence annotation

During manual recursive phylogenetic analyses of a gene family, gradually a model of gene evolution becomes apparent, despite contradicting topologies in the result trees (‘tree pool’). Some alternative topologies can be easily identified as artefacts, others remain questionable and require further study; each family has its own peculiarities in addition to the typical clade-specific characteristics. By interpreting the continuously accumulating phylogenetic gene trees, the at first complex model of gene evolution progressively turns into an easy to explain gene history. This model tree is then evaluated by annotating the quality and quantity of clade predictions from the gene trees. For each internode, we annotate the highest statistical support, the number of clade occurrence, the average clade support, and in addition we distinguish whether a clade was confirmed by a tree including all clade members or whether the clade is confirmed by a compatible clade with less members. By doing so, users can treat as unresolved (“soft polytomies”) nodes that are below the minimum required level of confidence. Confidence annotation is furthermore added in form of branch colors, indicating the strength of support or missing support. Latter gives a convenient survey already during the construction of SwissTree trees and can be used as guide for subsequent analyses. Vice versa, we annotate also the correctness of each pool tree given the model tree by shades of colors according to the gene tree’s branch support.

This procedure has been introduced to the SwissTree entries ST001, ST003, ST005, and ST018. The annotated model tree, pool trees, raw trees and MSAs are available at http://swisstree.vital-it.ch/gold_standard, the reference species tree is available at http://swisstree.vital-it.ch/species_tree.

#### SwissTree: maintenance and extendibility

Our framework makes the maintenance of a collection of deep gene histories feasible: upon new findings, for instance a change in the reference species tree, the updated gene tree models can be annotated with no further analysis based on the tree pool. Likewise, the tree pool can be augmented with trees reconstructed with alternative phylogenetic reconstruction methods or newly available data.

But what is more, the framework grants new options. Acquired - and generally conserved - molecular characteristics of genes, gene products or gene environments can be encoded as (partially or fully resolved) tree structures and added to the pool of gene trees, and their concordance or discordance with the model gene tree can be assessed. Examples of such complements are gene synteny, exon-intron structure, domain architecture, functional regions or sites.

**Fig. 6.**
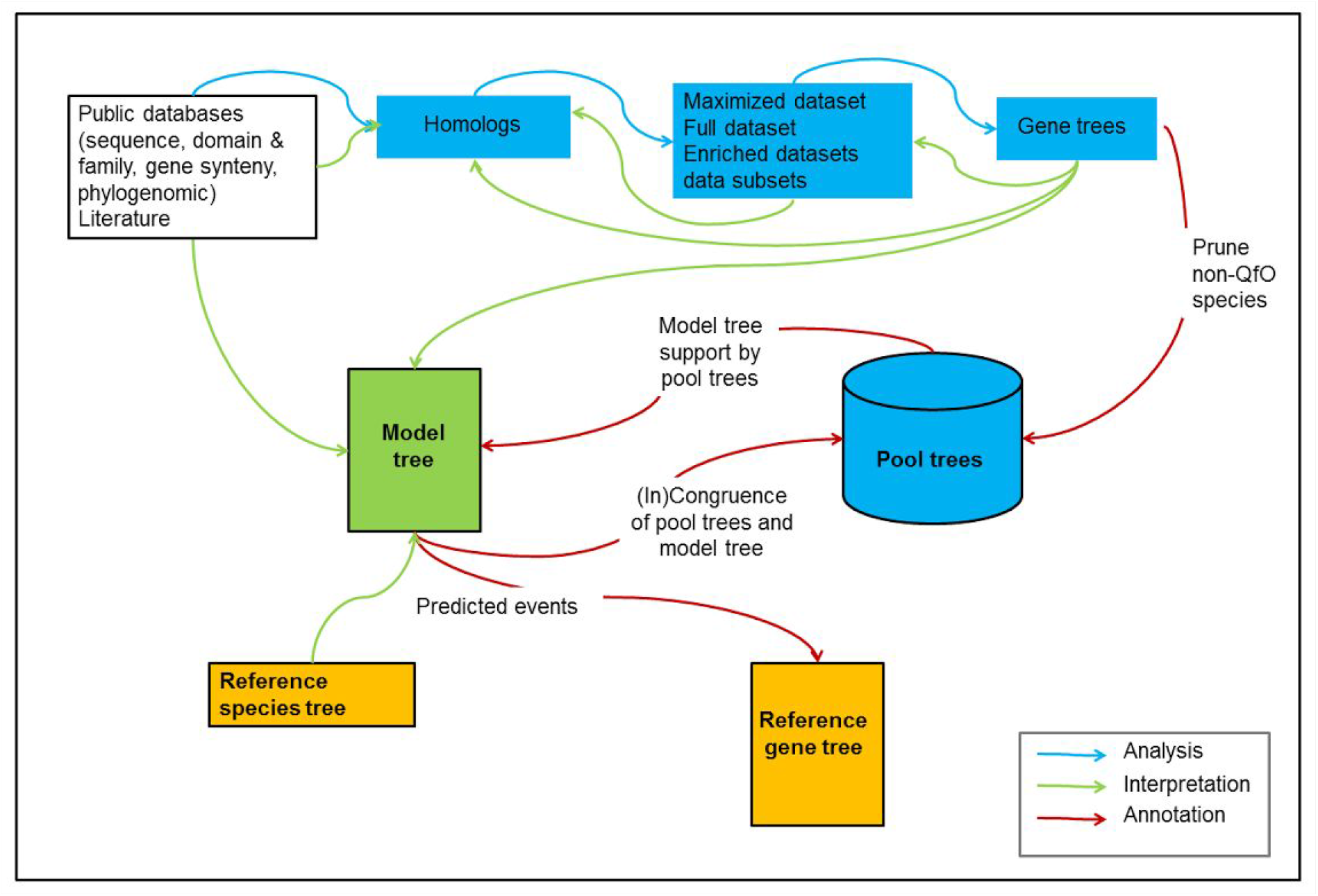
Conceptual overview of SwissTree construction.

### Case study: The APP gene family

Amyloid Precursor Proteins (APP) are cell-surface receptors involved in diverse functions including iron-export, cell adhesion, endocytosis, Notch signaling pathway inhibition, apoptosis. Well-studied is the impact of A4 subfamily members on neurons, because its cleavage can generate plaque-forming amyloid beta peptides in Alzheimer disease patients. Members of the gene family have been identified in the metazoan lineage. Typically, invertebrate genomes possess a single gene copy (APP), vertebrates three copies (A4, APLP1, APLP2), and ray-finned fishes an additional A4 gene copy (A4a, A4b). Four conserved domains are predicted in all family members analyzed here, three in the extracellular space (APP-N (PF02177), APP_Cu_bd (PF12924), APP_E2 (PF12925)) and one in the intracellular space (APP_amyloid (PF10515)). The vertebrate A4 and APLP2 subfamilies possess in addition a Kunitz domain (PF00014), and A4 subfamily members an extra beta-APP domain (PF03494). The placozoa *Trichoplax adhaerens* (UniProtKB mnemonic code: TRIAD) lacks a typical amyloid domain.

The phylogenetic analysis is performed on a set of 45 genes from Quest for Orthologs (QfO) reference species. The challenge is versatile. Protein sequences are highly conserved within mammalian subfamilies, resulting in a lack of phylogenetic signal particularly at the protein level for closely related species and in a strong mutational bias within the coding region of genes. By contrast, a lack of phylogenetic signal in invertebrate data is due to a low species density. Not to mention, subfamilies and clades evolve at different rates and multiple gene models are incomplete or erroneous. Based on the current dataset it therefore seems not feasible to generate an accurate gene family tree by means of a single analysis and indeed, none of the major phylogenomic databases at present show a conclusive gene phylogeny. However, by performing multiple problem-optimized analyses we can develop a model of gene evolution that is - in the end - in agreement with the reference species tree. Main clades, for instance, are analyzed with a dataset that is enriched with invertebrate genes and reduced for vertebrate genes. Subfamily phylogenies are inferred from nucleotide and amino acid sequence data by maximizing the alignment length and minimizing missing and ambiguous characters; latter have a strong impact on results when analyzing highly redundant data. Unstable or questionable nodes are re-evaluated in subsequent subclade-specific analyses. For instance in all the three subfamilies, the divergence order of primates, glires and laurasatherians is questionable and incongruence due to missing phylogenetic signal can be expected because of short time spans between the two speciation events. Because the species tree-consistent topology is one of the two alternative topologies and we observe no evidence for an evolutionary event other than speciation, the model tree is concordant with the species tree at this node, annotated with the corresponding bootstrap support. Figure 7 shows an example of a phylogenetic tree that was optimized for the study of early vertebrate divergence patterns. During the analysis we start with enlarged datasets and strive to stepwise reduce redundancies while the topology remains consistent. This way, we can also remove taxa for which species phylogenies are inconsistent, thus making an interpretation of the tree difficult. The model gene tree for the APP family is shown in Figure 7 along with one of the pool trees and the corresponding raw tree. The annotated model tree, pool trees and raw trees can be explored at http://swisstree.vital-it.ch/ST018.

**Figure 7.**
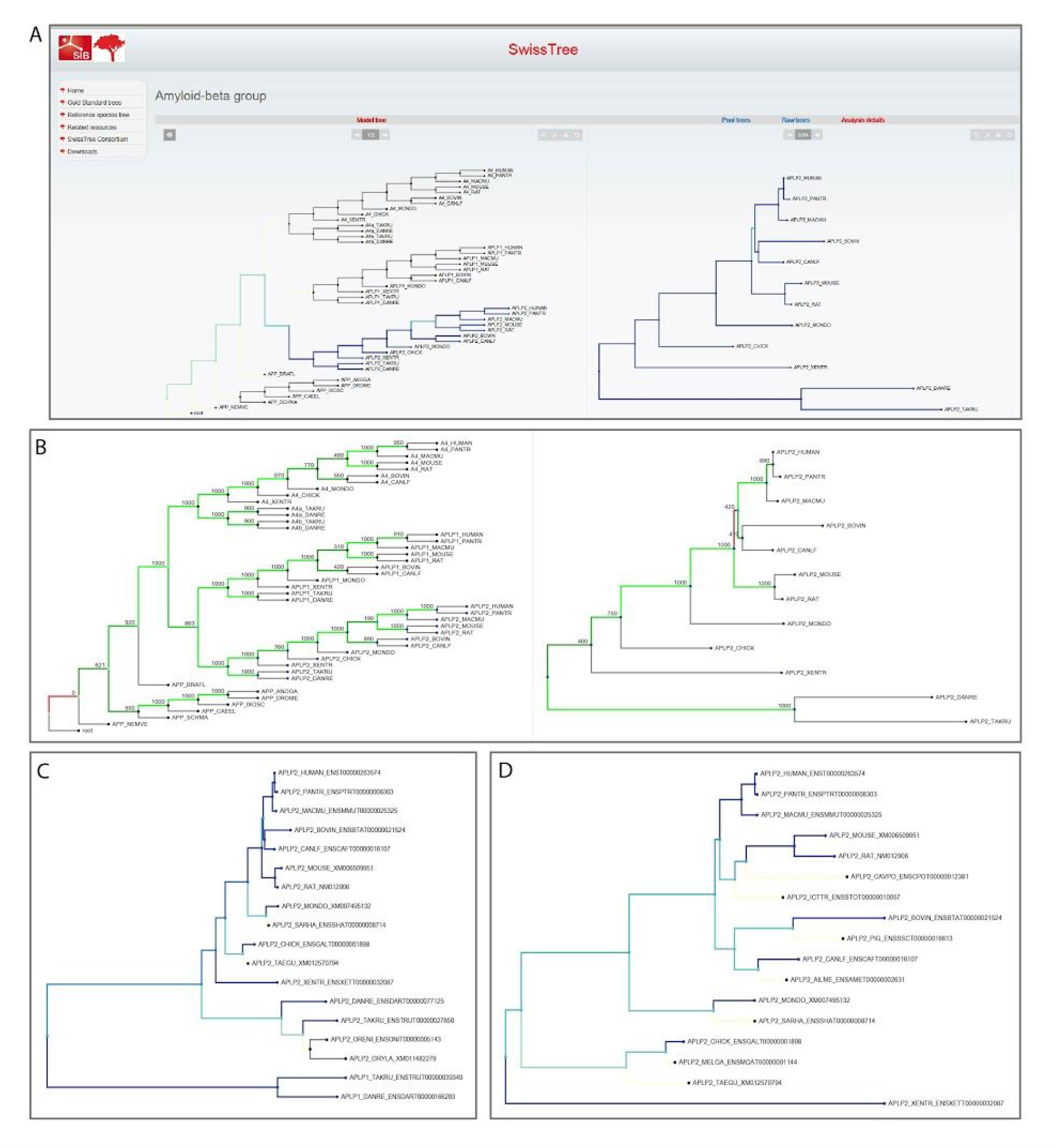
SwissTree model gene tree for the APP gene family (ST018). A. Comparison of the model tree (left) and one of the pool trees (tree 6) for subfamily APLP2 (right), visualized using phylo.io with tree comparison setting; tree concordance is color-coded dark blue, discordance light blue, missing clades are shown in white, missing subclades in grey. The analyzed subfamily APLP2 stands out in blue. B. Same as A, ‘maximum support’ settings; the model tree is annotated and color-coded according to the maximum level of clade support by the pool trees, the pool tree is color-coded for model tree concordance and shows the level of bootstrap support. Green indicates topological concordance, the color shade relates to the level of support; red indicates topological discordance. The outgroup branch of the model tree is always shown in red, because outgroups are removed from the pool trees prior to tree reconciliation. In the pool tree the primate/laurasatherian clade is not confirmed. C. Raw tree corresponding to the pool tree shown in Figure A and B, color-coded by topological concordance with the model tree. Raw trees are phylogenetic gene trees used to generate pool trees by pruning supplementary taxa and subsequent annotation. The raw tree shows additional taxa in for early vertebrates (white and grey branches), which was the focus of interest in this analysis. D. Raw tree generated with a focus on the mammalian clade. The enrichment of the dataset with mammalian genes (white and grey branches) results in a tree topology that is in concordance with the species tree.

### Large-scale study. Synapomorphic signal, tree inconsistency and taxa sampling (*Boreoeutheria*)

Phylogenetic analyses of boreoeutherians lead to controversial phylogenies for primates, glires and laurasatherians, supporting either a primate glires sister clade (e.g. [15], [16][17] or a primate laurasatherian sister clade (e.g. [18], [19], [20]). QfO datasets include seven boreoeutherian proteomes, three primates (human, chimpanzee, macaque), two glires (mouse, rat) and two laurasatherians (bovin, dog) (abbreviated by 7(3/2/2) in Fig. 8). In a similar way than in the simulation study above, we analyze MSAs from 374 1:1 orthologous gene families for synapomorphic characters. Results clearly favor a primate laurasatherian sister clade with glires as outgroup (Fig. 8A). Because the dataset is small and furthermore redundant for glires, caution is advised for the interpretation of these results. More importantly, this data is used for benchmarking purposes and we explore topologies of corresponding phylogenetic trees from a phylogenomic database that does not take into account species trees for tree reconstruction. In agreement with the phylogenetic signals observed in the MSAs, we find indeed highest clade occurrence frequencies for the sister clade primates/laurasatherians (Fig. 8B). In addition, we observe within gene families tree inconsistency that corresponds to the fraction of observed synapomorphic and contradicting signal. This result suggests that the QfO benchmarking dataset of seven relevant proteomes could be indeed too small for a rational inference of the boreoeutherian phylogeny, in which case results should differ when analyzing enlarged datasets. To test this, we perform the same MSA analysis on a dataset of 3526 mammalian families including at least six members of each clade of interest and at least two out of six selected outgroup members. From this dataset, three subsets are generated: 1) the seven relevant species of the QfO datasets (same as above; 7(3/2/2)), 2) all six selected species for each of the three clades (18(6/6/6)), and 3) two diverse species for each of the three clades (6(2/2/2)) (Fig. 8C-E). Results suggest that MSAs from the large dataset and from the small dataset of diverse species contain more synapomorphic signal for the established human mouse sister clade (Euarchontoglires) than for alternative tree topologies. Now going back to findings from previous publications, it is striking that it is especially the innovative large-scale studies that suggest the primate laurasatherian sister clade, but because such analyses are CPU-time intensive and because of the small number of genomes with a high quality gene prediction one to two decades ago, the number of genomes used was often low. Rerunning the same approaches with data from more diverse species, might well result in consistent topologies with similar support.

**Figure 8.**
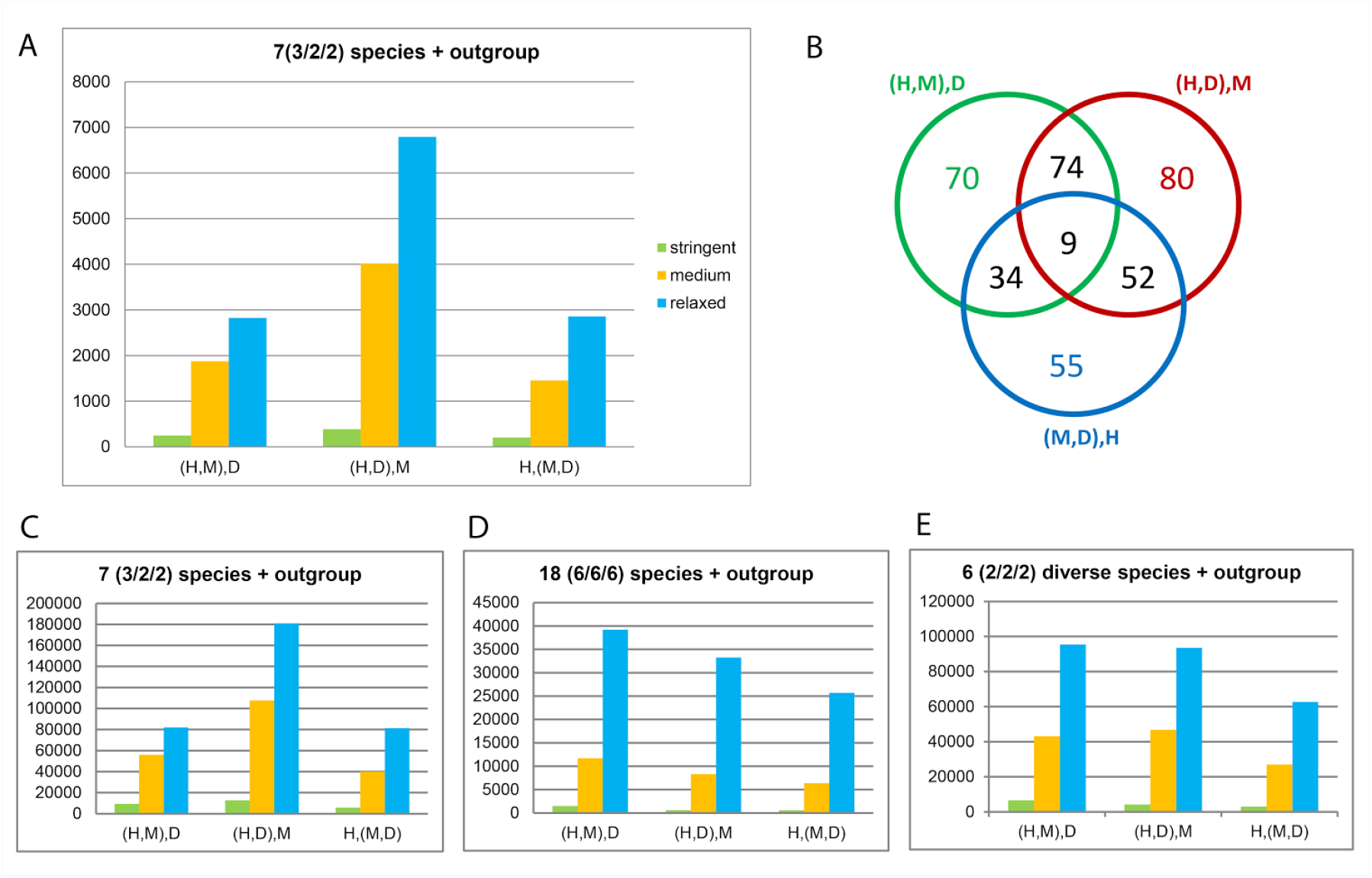
Quantitative assessment of synapomorphic signal for boreoeutherian sister clades. A. Number of synapomorphic positions from MSAs of 374 one-to-one ortholog gene families for tree topologies with three different boreoeutherian sister clades primates/glires (H,M), primates/laurasatherian (H,D) and glires/laurasatherians (M,D); tree topologies are exemplified for Human (H), Mouse (M), Dog (D); for more detail, see text and cmp. Fig. 4. B. Venn diagram depicting the observed tree heterogeneity in the 374 gene families with 2-17 trees obtained from different datasets of the QfO proteomes. The largest fraction of families supports a primate/laurasatherian sister clade, tree heterogeneity is observed in 45.2% of the gene families. C-E. Results from a dataset of 25,272 MSAs. C. Same taxon composition as in A; results confirm findings in A. D. Enlarged dataset with six diverse species for each of the three clades; the largest fraction of signal supports the primates/glires sister clade; E. Dataset with only two, but diverse, representative species from each clade.

### Assessment of orthology predictions with SwissTree

Benchmarking results with SwissTree are obtained from the Orthology Benchmarking Webservice (http://orthology.benchmarkservice.org; [8] for 15 sets of orthology predictions from important methods and databases: the pairwise methods InParanoid [21] and a higher confidence subset named “InParanoid core”, OMA pairs [22], Reciprocal Best Hits (RBH) [23] and Reciprocal Smallest Distance (RSD) [24], the group-wise (graph-based) methods EggNOG [25], Hieranoid [26], OMA groups [22], OMA GETHOGs [27] and OrthoInspector [28], and the tree-based methods Ensembl Compara [29], PANTHER [30], a subset of PANTHER named PANTHER_LDO (Least Diverged Orthologs), and PhylomeDB [31], as well as the meta-method MetaPhOrs [32] (for details, see [8]). A survey on the correctness and completeness of predicted orthologs from the benchmarking dataset reveals a precision-recall trade-off for all methods from all database concepts (pairs, groups, trees, meta), particularly when higher confidence predictions are provided in a subset (InParanoid_core, PANTHER_LDO) (Fig. 9A, Supporting Information S3, Table S3-1).

Three factors vitally affect the identification of orthologs: the sequence length, the evolutionary distance between proteins and the complexity of gene families. The sequence length confines the region within which phylogenetic signals can occur and logically, long sequences can capture more signal than short ones. Indeed, the distribution of sequence lengths for correct, missing and incorrect predictions show the trend for higher correctness at long sequence length (mean: 413.99 aa), and more false predictions (FP, FN) at shorter sequence length (mean: 333,79 aa) (Fig. 9B, Supporting Information S3, Fig. S3-1). Another challenge for orthology prediction is the difference in sequence length of gene pairs, which can occur naturally or as an analysis artefact. On the one hand, sequence search and alignment strategies can be more difficult than for equally long sequences, on the other hand it is a fundamental decision whether to consider also partial sequence homologies or not. Phylogenomic databases deal with this issue by setting cut-offs for minimal sequence length differences or overlaps in the analysis procedure, which can furthermore be combined with a minimum sequence identity or similarity score. Thereby, a stringent cut-off increases the number of missed predictions (FN), a relaxed cut-off the number of incorrect predictions (FP). It is thus an important feature of a database’s strategy and an essential criterion for users in search for a dataset suitable for a specific research question. Figure 9C (Supporting Information S3, Fig. S3-2) illustrates the correlation between the sequence length differences of gene pairs and the prediction accuracy: on average the length of correctly predicted orthologs is more similar (mean: 39.16 aa) than the length of incorrectly predicted orthologs (mean: 145.42 aa). For pairwise methods, particularly RBH and RSD, largest sequence differences are observed for incorrect predictions (mean: 201.30 aa), graph-based methods show strongest length differences either for missing orthologs or incorrect predictions, and tree-based methods possess a mostly balanced distribution of sequence length differences for missing and incorrect predictions. It is especially the orthologs from distant species that largely differ in size, so that results might be influenced from enlarged evolutionary distances. By testing the impact of evolutionary distances on the analysis results, we observe indeed similar trends for correct and incorrect predictions (substitutions per site; mean for TP: 1.21; mean for FP/FN: 2.13) (Fig. 9D, Supporting Information S3, Fig. S3-3).

Gene families are difficult to analyze when genes duplicate successively within a short time range and when paralogs evolve at different rates. We grouped families into two categories of complexity, small (S) and large (L), according to the number of paralogs. Taken all results from databases together, the fraction of false positives is considerably higher in large families (1.05%) than in small families (0.25%). By contrast, the fraction of missing orthologs in large families is only about half of that from small families (24,26% versus 12.25% in large families) for the set of SwissTree families.

**Figure 9.**
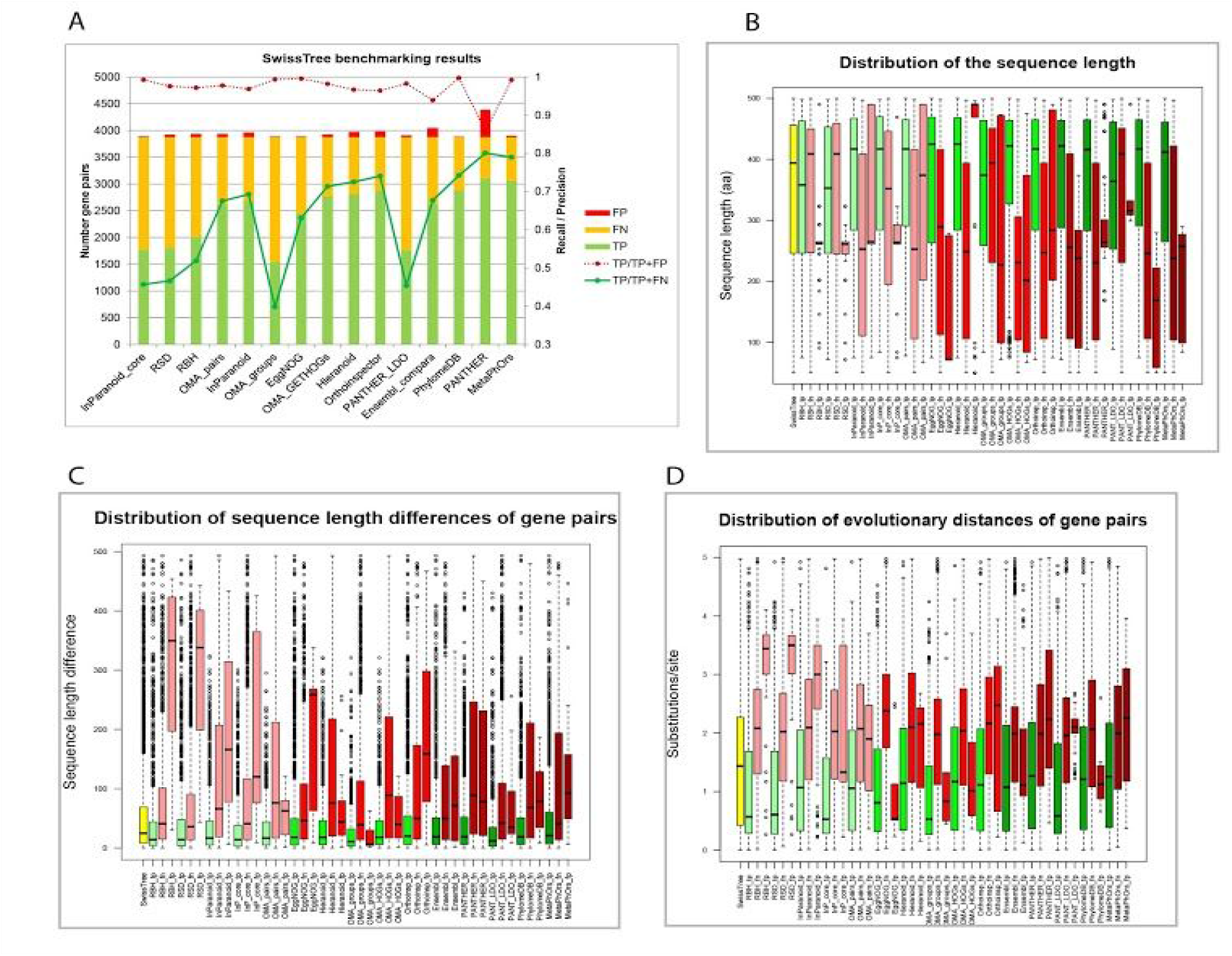
Survey on SwissTree benchmarking results. A. Correct (TP) and incorrect (FP, FN) orthology predictions for 15 approaches, sorted by increasing number of true positives for each orthology prediction strategy in the order pairs, groups, trees, meta. B.-D. Distribution of sequence lengths (B), sequence length differences between genes of gene pairs (C) and evolutionary distances (D) for correct (green) and incorrect predictions (red) in the order pairs (bright-colored boxes), groups (medium-colored), trees (darkish), meta (darkest). The first, yellow box shows the distribution of values for orthologs predicted by SwissTree. Figures are available in Supporting Information S3.

### Outlook - Automation perspective of the SwissTree procedure

In this study we demonstrate a new approach that could serve as a basis of sustainable representative gene trees. Attentive taxon sampling optimized for the analysis of large extended datasets, individual clades and subclades guaranties a comprehensive data exploration. The thereby generated gene tree heterogeneity conduces to the identification of analysis artefacts and missing or contradicting phylogenetic signal. In addition, potential cases of discordant evolution (e.g. introgression, ILS) stand out. Annotation, maintenance and extensibility of model gene trees are part of the SwissTree concept.

During the development, analyses were performed manually. Without bearing automation in mind, optimization steps are at the risk of introducing bias. It is therefore eligible to progressively automate the analysis procedure, whereby an elaborated taxon sampling occupies a central position in view of a massive amount of genomic data. The second suspenseful topic concerns the construction of preliminary model gene trees. With some experience we might learn the rules to generate assumed models automatically for a large part of genes and to subsequently evaluate concordance with the phylogenetic gene trees. By doing so, human time will be freed to focus on unconfirmed branches in preliminary model gene trees.

## Material and Methods

### Simulation data

A simulation dataset of 1000 protein-coding, 1:1 orthologs for 100 taxa was constructed with ALF, with speciation events occurring according to a ToL of 1038 species from the OMA project (server version 3-Aug-2015 at http://alfsim.org; [33]) from a generated root genome (parameter settings: realseed false, minGeneLength=240, gammaLengthDist=2.4/133.8, blocksize=3, treeType=TolSample, scaleTree=false, substModels=CPAM [34], indelModel=(0.0003, ZIPF, 1.821,50), rateVarModel=(gamma,5,0.01,1), targetFreqs=random, amongGeneDistr=(gamma,1), time scale PAM). The amount of possible phylogenetic information that a sequence can carry is limited by the sequence length. The minimal gene length of the ancestral genome is set to 240 nucleotides; the simulated genes evolve to a length between 177 and 4083 nucleotides, 1071 on average (Fig. 10). The result includes the species tree and the MSA as well as the phylogenetic tree for each gene family. The distribution of the gene lengths are given below.

**Fig. 10.**
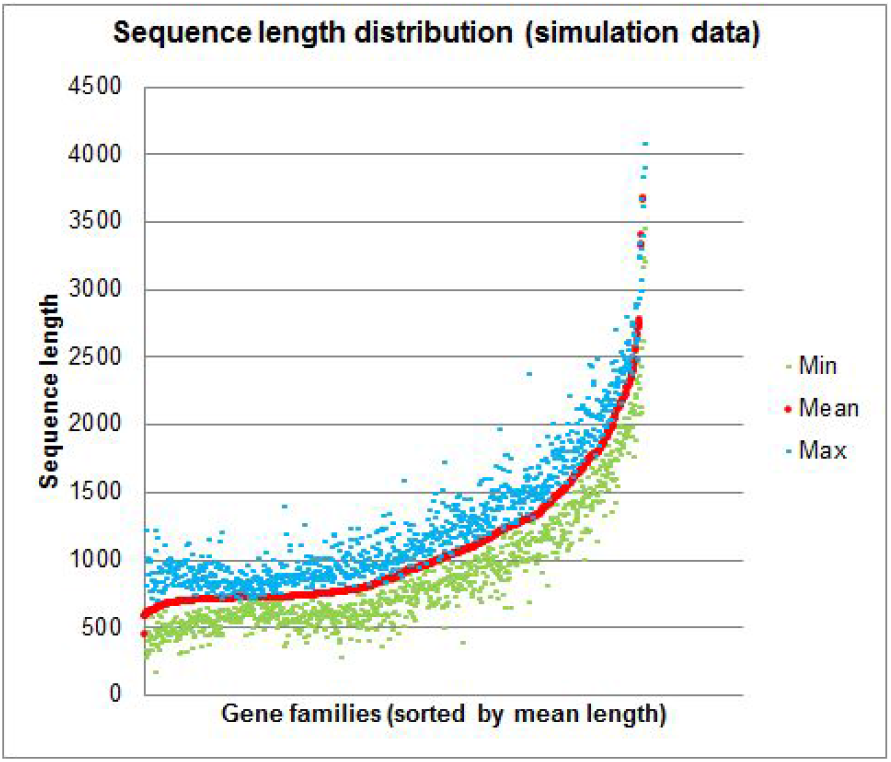
Scatterplot depicting the range of nucleotide sequence lengths for each of the 1000 simulated gene families.

From the full simulated dataset, we generated six data subsets: 1) a balanced set of 30 taxa, obtained by successively pruning taxa with long branches and small clades with short internodes from the 100 species tree (b30); 2) a balanced set of 10 taxa, obtained by successively pruning taxa with long branches and small clades with short internodes from the species tree of dataset b30 (b10); 3) 30 randomly selected taxa from the 100 taxa (r30); 4) 10 randomly selected taxa from 100 taxa (r10); 5) a subclade of 30 taxa of the 100 species tree (a30); 6) a subclade of 10 taxa of the 30 species tree (a10). The full dataset is named a100, and we further distinguish a100/aa for protein sequence data, and a100/codon and a100/dna for nucleotide sequence data. Taxon names and the known tree topology for each dataset are given in the Supporting Information S1.

### Phylogenetic analysis of the simulation data

Alignments for each dataset are extracted from the true alignments and gap positions with less than 10% characters are removed from the dataset in order to save computing time. Trees are reconstructed with codon-PhyML using the model HKY85+F+G(4)+I for nucleotide substitution, GY+W+K+F F3x4 for codon substitution, and WAG+F+G(4)+I for amino acid substitution. These models are likely over-parameterized in most cases. In the ML and BI frameworks and for nucleotide substitution models, overfitting was found to be robust on tree topologies, but can affect branch length estimates, branch support and the overall likelihood and posterior probability of a tree, respectively [35],[36]. The tree topology search was performed with NNI and SPR (‘Both’), except for the codon-based analysis for which we applied only NNI to minimize a possible advantage for this approach over others because the simulation data evolved under a codon-based model. Parameters are optimized for tree topology, branch length and substitution rate. For each dataset 1000 maximum likelihood (ML) trees were inferred under the model of nucleotide, codon and amino acid substitution, resulting in 21000 trees.

For the comparison of clade recurrence values and aLRT-SH support from complete proteomes versus single datasets, we selected from the list of all genes sorted by alignment length every 50th alignment, starting at position 50. The known alignments of the 20 genes were obtained for datasets a30, b30 and r30 and a rapid bootstrap analysis was performed from 1000 replicates using RAxML (model: GAMMAIWAGF).

### Analysis of tree correctness

The concordance of reconstructed gene trees with the corresponding species trees are measured using Perl scripts; the fraction of correct and incorrect trees as well as the fraction of correct and incorrect clades are obtained directly from the annotated sequence trees. Results are visualized and compared with R (3.0.3) and Microsoft Excel (Office14). To assess characteristics of balanced trees, we plot for dataset a100/aa confidence values from correct clade predictions (clade occurrence frequency, branch support (aLRT-SH), maximum branch support (aLRT-SH) against the corresponding internode length from the species tree (PAM distance). Confidence of corresponding correct clades from different datasets are obtained by mapping compatible clades between the a100 species tree and the species trees of the six sub-datasets.

### Analysis of synapomorphic sites in the known MSAs of the simulation data

Amino acids assumed to be a shared derived character of a sister clade is denoted ‘synapomorphic’. The number of synapomorphic sites in the MSA is determined in three modes. In the stringent mode, the two sister clades share a conserved amino acid in an MSA position, and the two outgroups share another conserved amino acid, whereas in the medium mode, the two outgroups to the sister clade can have any amino acid except the one shared by the sister clade. In the relaxed mode, the two sister clades share at least one amino acid in an MSA position, and the two outgroups to the sister clade can possess any amino acid except the one shared by the sister clade. The number of synapomorphic sites is determined for two clades (63, 87) for the true and 2 alternative divergence patterns of the subclades (Fig. 4A) from the 1000 MSAs of the amino acid sequences from all simulated datasets. In addition, we count the number of parsimonious non-informative sites according to our criteria (considering only identity, not similarity) and sites that are not considered (gaps: missing positions in at least one clades or in the outgroup).

### Consensus tree construction and ranking strategies

Consensus trees are constructed with Perl scripts, following a Nelson-like combinable-component approach for unrooted trees. From a set of unrooted gene trees, tree split information is summarized in a matrix to ease the mapping of corresponding splits and to collect clade confidence information such as clade occurrence frequency, maximum branch support and the sum of branch support for each clade, which is used to generate ordered clade lists for consensus tree construction. As ranking criteria, we use clade occurrence frequencies as well as ‘mean2’ which is the arithmetic mean of the clade support considering the clade occurrence frequency in that the clade support for not predicted true clades is set to zero. Consensus trees are constructed for the three trees for each gene (DNA, codon, aa; dataset), for the three gene trees pruned from the a100 gene trees (DNA, codon, aa; pruned), as well as for all six gene trees (DNA, codon aa; dataset and pruned). Trees of the largest datasets (a100/dna, a100/codon, a100/aa) were pruned to the size of the six data subsets using the Newick utilities [37].

### Phylogenetic analysis of the APP gene family

The phylogenetic analysis is performed on gene data from 19 out of 21 metazoan Quest for Orthologs (QfO) reference species (04_2016; http://www.ebi.ac.uk/reference_proteomes); the two metazoans not included are *Ciona intestinalis* (vase tunicate) with no predicted APP gene family members, and *Ornithorhynchus anatinus* (platypus) with largely incomplete data. Transcript and protein sequence data is obtained from UniProt [38], ENSEMBL [39], and NCBI [40]. Transcripts are translated into amino acid sequences, aligned (T-coffee: Expresso mode [41]; MAFFT: INSi-E or INSi-G mode [42], dependent on the domain composition of the dataset) and subsequently explored and edited using Jalview [43] or MEGA (v6.06) [44]. The nucleotide sequences are mapped to the alignment, and the data model is selected manually in order to maximize homologous sequence regions and to remove ambiguous positions. Best fit models are determined with Prottest (v3.4) [45] and Mega6 (v6.06) and ML trees reconstructed using codonPhyml and Mega6. The resulting trees are inspected with Archaeopteryx (https://sites.google.com/site/cmzmasek/home/software/archaeopteryx) or phylo.io [46]. If phylogenetic gene trees are incongruent with the SwissTree reference species tree model or weakly supported, the stability of tree topologies is revised by subclade analyses with a higher species density. If the new gene tree is concordant with the species tree, the tree is pruned to the species set of interest (using the NEWICK Utilities or manually with Archaeopteryx) and added to the gene tree pool. The analysis of the family and subfamily results in x trees. Subsequent to the analysis of ambiguous clades, a model of gene family phylogeny is generated manually. The gene tree support - occurrence frequency, highest support and mean support - is mapped to all nodes to the family tree model (extended Newick format) using Perl scripts; likewise, congruence with the gene tree model is annotated in all gene trees.

### Analysis of synapomorphic sites in MSAs and tree inconsistency within gene families

The PhylomeDB/QfO benchmarking reference dataset 2013 comprises 186282 trees, from which we remove exact tree duplicates (30753 trees), prune prokaryotic genes from all trees (991613 OTUs; prokaryotic data is not suitable for this analysis because of extensive HGT and the lack of a confident species tree) and discard trees with less than 4 OTUs (12192 trees). Subsequently, trees with two or more genes from the same species (132584 trees with intra-species gene copies) are filtered. Gene trees that share at least one gene are grouped into 2262 families and filtered for intra-species gene copies across trees (349 families), resulting in 1913 1:1 ortholog gene families with at least two gene trees. This dataset is used to study tree inconsistency within gene families. Model trees are generated for each family by pruning the species tree to the relevant set of family members. Family tree pools and model trees are reconciled, annotated and evaluated as described above. In order to calculate the maximal possible support for each node, we perform the same analysis with trees obtained by pruning the species tree to the corresponding set of family members for each pool tree. To determine support for different models of mammalian evolution, we select from the set of 1913 families those with the relevant species in the QFO benchmarking dataset, namely three primates *Homo sapiens, Pan troglodytes, Macaca mulatta*), two glires (*Mus musculus, Rattus norvegicus*) and two laurasatherian (*Bos taurus, Canis lupus familiaris*), resulting in 374 families. Because models are rooted, there is no need to include outgroups. Two alternative model trees are generated with the sister clades primates/laurasatherians and laurasatherians/glires. The analysis is performed as described above. From the corresponding MSAs of the human phylome, we determine the number of synapomorphic sites for the true tree and the 2 alternative trees around an internode in the three modes for the 374 families of the QfO benchmarking dataset.

From a dataset of 26252 mammalian MSAs provided by eggNOG (maNOG; release 4.5.1), we select 3526 MSAs that contain a single (assumed 1:1 orthologs) of six primates (*Homo sapiens, Pan troglodytes, Pongo abelii, Macaca mulatta, Callithrix jacchus, Microcebus murinus*), glires (*Cavia porcellus, Dipodomys ordii, Ictidomys tridecemlineatus, Mus musculus, Oryctolagus cuniculus, Rattus norvegicus*), and laurasatherians (*Bos taurus, Canis lupus familiaris, Equus caballus, Myotis lucifugus, Sus scrofa, Tursiops truncatus*) as well as at least two outgroup members (*Loxodonta africana, Dasypus novemcinctus*, *Monodelphis domestica, Notamacropus eugenii, Ornithorhynchus anatinus, Sarcophilus harrisii*). The number of synapomorphic positions is determined for three datasets: 1) the seven species of the QfO benchmarking dataset 2013 (same as above), 2) for a dataset of 6 species per clade and for a dataset of two diverse species per clade (*Homo sapiens, Microcebus murinus, Mus musculus, Oryctolagus cuniculus*, *Bos taurus, Myotis lucifugus*).

### Analysis of the SwissTree benchmarking results

Benchmarking results for SwissTree were obtained from the orthology benchmarking service (http://orthology.benchmarkservice.org) in form of lists including the unique identifiers for each gene of a gene pair as well as the prediction result (True Positive (TP), True Negative (TN), False Positive (FP), False Negative (FN)). From the QfO benchmarking proteomes we determined the length of each sequence and calculated the sequence length difference of gene pairs. The evolutionary distance of gene pairs was estimated from the multiple sequence alignment of each family using MEGA6.6. Finally we classified gene families in three categories of complexity according to the number of paralogs. Database concept-specific prediction results were determined by calculating Venn diagrams at http://bioinformatics.psb.ugent.be/webtools/Venn.

### Visualization

Graphs were generated using R and MS-Excel. Phylogenetic trees are visualized with phylo.io.

## Acknowledgements

This work was supported by the Swiss State Secretariat for Education, Research and Innovation (SERI) funding (BB). CD acknowledges support by Swiss National Science Foundation grant 150654 and by the Swiss Institute of Bioinformatics for the tree visualisation tool phylo.io.

## References

1. Nabhan AR, Sarkar IN. The impact of taxon sampling on phylogenetic inference: a review of two decades of controversy. Brief Bioinform. 2012; 13: 122–134.

2. Szöllősi GJ, Tannier E, Daubin V, Boussau B. The inference of gene trees with species trees. Syst Biol. 2015; 64: e42–62.

3. Sonnhammer ELL, Gabaldón T, Sousa da Silva AW, Martin M, Robinson-Rechavi M, Boeckmann B, et al. Big data and other challenges in the quest for orthologs. Bioinformatics. 2014; 30: 2993–2998.

4. Trachana K, Larsson TA, Powell S, Chen W-H, Doerks T, Muller J, et al. Orthology prediction methods: a quality assessment using curated protein families. Bioessays. 2011; 33: 769–780.

5. Schreiber F, Patricio M, Muffato M, Pignatelli M, Bateman A. TreeFam v9: a new website, more species and orthology-on-the-fly. Nucleic Acids Res. 2014; 42: D922–5.

6. Boeckmann B, Marcet-Houben M, Rees JA, Forslund K, Huerta-Cepas J, Muffato M, et al. Quest for Orthologs Entails Quest for Tree of Life: In Search of the Gene Stream. Genome Biol Evol. 2015; 7: 1988–1999.

7. Boeckmann B, Robinson-Rechavi M, Xenarios I, Dessimoz C. Conceptual framework and pilot study to benchmark phylogenomic databases based on reference gene trees. Brief Bioinform. 2011; 12: 423–435.

8. Altenhoff AM, Boeckmann B, Capella-Gutierrez S, Dalquen DA, DeLuca T, Forslund K, et al. Standardized benchmarking in the quest for orthologs. Nat Methods. 2016; 13: 425–430.

9. Rasmussen MD, Kellis M. Unified modeling of gene duplication, loss, and coalescence using a locus tree. Genome Res. 2012; 22: 755–765.

10. Efron B, Halloran E, Holmes S. Bootstrap confidence levels for phylogenetic trees. Proc Natl Acad Sci U S A. 1996; 93: 7085–7090.

11. Anisimova M, Gil M, Dufayard J-F, Dessimoz C, Gascuel O. Survey of Branch Support Methods Demonstrates Accuracy, Power, and Robustness of Fast Likelihood-based Approximation Schemes. Syst Biol. 2011; 60: 685–699.

12. Lecointre G, Philippe H, Vân Lê HL, Le Guyader H. Species sampling has a major impact on phylogenetic inference. Mol Phylogenet Evol. 1993; 2: 205–224.

13. Wägele JW, Mayer C. Visualizing differences in phylogenetic information content of alignments and distinction of three classes of long-branch effects. BMC Evol Biol. 2007; 7: 147.

14. Dessimoz C, Boeckmann B, Roth ACJ, Gonnet GH. Detecting non-orthology in the COGs database and other approaches grouping orthologs using genome-specific best hits. Nucleic Acids Res. 2006; 34: 3309–3316.

15. Murphy WJ, Eizirik E, Johnson WE, Zhang YP, Ryder OA, O’Brien SJ. Molecular phylogenetics and the origins of placental mammals. Nature. 2001; 409: 614–618.

16. Madsen O, Scally M, Douady CJ, Kao DJ, DeBry RW, Adkins R, et al. Parallel adaptive radiations in two major clades of placental mammals. Nature. 2001; 409: 610–614.

17. Song S, Liu L, Edwards SV, Wu S. Resolving conflict in eutherian mammal phylogeny using phylogenomics and the multispecies coalescent model. Proc Natl Acad Sci U S A. 2012; 109: 14942–14947.

18. Cannarozzi G, Schneider A, Gonnet G. A phylogenomic study of human, dog, and mouse. PLoS Comput Biol. 2007; 3: e2.

19. Huttley GA, Wakefield MJ, Easteal S. Rates of genome evolution and branching order from whole genome analysis. Mol Biol Evol. 2007; 24: 1722–1730.

20. Lin Y, Rajan V, Moret BM. Bootstrapping phylogenies inferred from rearrangement data. Algorithms Mol Biol. 2012; 7: 21.

21. Ostlund G, Schmitt T, Forslund K, Köstler T, Messina DN, Roopra S, et al. InParanoid 7: new algorithms and tools for eukaryotic orthology analysis. Nucleic Acids Res. 2010; 38: D196–203.

22. Altenhoff AM, Škunca N, Glover N, Train C-M, Sueki A, Piližota I, et al. The OMA orthology database in 2015: function predictions, better plant support, synteny view and other improvements. Nucleic Acids Res. 2015; 43: D240–9.

23. Overbeek R, Fonstein M, D’Souza M, Pusch GD, Maltsev N. The use of gene clusters to infer functional coupling. Proc Natl Acad Sci U S A. 1999; 96: 2896–2901.

24. Wall DP, Fraser HB, Hirsh AE. Detecting putative orthologs. Bioinformatics. 2003; 19: 1710–1711.

25. Powell S, Forslund K, Szklarczyk D, Trachana K, Roth A, Huerta-Cepas J, et al. eggNOG v4.0: nested orthology inference across 3686 organisms. Nucleic Acids Res. 2014; 42: D231–9.

26. Schreiber F, Sonnhammer ELL. Hieranoid: hierarchical orthology inference. J Mol Biol. 2013; 425: 2072–2081.

27. Altenhoff AM, Gil M, Gonnet GH, Dessimoz C. Inferring hierarchical orthologous groups from orthologous gene pairs. PLoS One. 2013; 8: e53786.

28. Linard B, Allot A, Schneider R, Morel C, Ripp R, Bigler M, et al. OrthoInspector 2.0: Software and database updates. Bioinformatics. 2015; 31: 447–448.

29. Vilella AJ, Severin J, Ureta-Vidal A, Heng L, Durbin R, Birney E. EnsemblCompara GeneTrees: Complete, duplication-aware phylogenetic trees in vertebrates. Genome Res. 2009; 19: 327–335.

30. Mi H, Muruganujan A, Thomas PD. PANTHER in 2013: modeling the evolution of gene function, and other gene attributes, in the context of phylogenetic trees. Nucleic Acids Res. 2013; 41: D377–86.

31. Huerta-Cepas J, Capella-Gutiérrez S, Pryszcz LP, Marcet-Houben M, Gabaldón T. PhylomeDB v4: zooming into the plurality of evolutionary histories of a genome. Nucleic Acids Res. 2014; 42: D897–902.

32. Pryszcz LP, Huerta-Cepas J, Gabaldón T. MetaPhOrs: orthology and paralogy predictions from multiple phylogenetic evidence using a consistency-based confidence score. Nucleic Acids Res. 2011; 39: e32.

33. Dalquen DA, Anisimova M, Gonnet GH, Dessimoz C. ALF––a simulation framework for genome evolution. Mol Biol Evol. 2012; 29: 1115–1123.

34. Schneider A, Cannarozzi GM, Gonnet GH. Empirical codon substitution matrix. BMC Bioinformatics. 2005; 6: 134.

35. Sullivan J, Swofford DL. Should we use model-based methods for phylogenetic inference when we know that assumptions about among-site rate variation and nucleotide substitution pattern are violated? Syst Biol. 2001; 50: 723–729.

36. Huelsenbeck J, Rannala B. Frequentist properties of Bayesian posterior probabilities of phylogenetic trees under simple and complex substitution models. Syst Biol. 2004; 53: 904–913.

37. Junier T, Zdobnov EM. The Newick utilities: high-throughput phylogenetic tree processing in the UNIX shell. Bioinformatics. 2010; 26: 1669–1670.

38. The UniProt Consortium. UniProt: the universal protein knowledgebase. Nucleic Acids Res. 2017; 45: D158–D169.

39. Aken BL, Achuthan P, Akanni W, Amode MR, Bernsdorff F, Bhai J, et al. Ensembl 2017. Nucleic Acids Res. 2017; 45: D635–D642.

40. O’Leary NA, Wright MW, Brister JR, Ciufo S, Haddad D, McVeigh R, et al. Reference sequence (RefSeq) database at NCBI: current status, taxonomic expansion, and functional annotation. Nucleic Acids Res. 2016; 44: D733–45.

41. Di Tommaso P, Moretti S, Xenarios I, Orobitg M, Montanyola A, Chang J-M, et al. T-Coffee: a web server for the multiple sequence alignment of protein and RNA sequences using structural information and homology extension. Nucleic Acids Res. 2011; 39: W13–7.

42. Katoh K, Standley DM. A simple method to control over-alignment in the MAFFT multiple sequence alignment program. Bioinformatics. 2016; 32: 1933–1942.

43. Waterhouse AM, Procter JB, Martin DMA, Clamp M, Barton GJ. Jalview Version 2––a multiple sequence alignment editor and analysis workbench. Bioinformatics. 2009; 25: 1189–1191.

44. Kumar S, Stecher G, Tamura K. MEGA7: Molecular Evolutionary Genetics Analysis Version 7.0 for Bigger Datasets. Mol Biol Evol. 2016; 33: 1870–1874.

45. Darriba D, Taboada GL, Doallo R, Posada D. ProtTest 3: fast selection of best-fit models of protein evolution. Bioinformatics. 2011; 27: 1164–1165.

46. Robinson O, Dylus D, Dessimoz C. Phylo.io: Interactive Viewing and Comparison of Large Phylogenetic Trees on the Web. Mol Biol Evol. 2016; 33: 2163–2166.

